# Establishing farm dust as a useful viral metagenomic surveillance matrix

**DOI:** 10.1101/2021.03.09.434704

**Authors:** Kirsty T. T. Kwok, Myrna M. T. de Rooij, Aniek B. Messink, Inge M. Wouters, Lidwien A. M. Smit, Matthew Cotten, Dick J.J. Heederik, Marion P. G. Koopmans, My V. T. Phan

## Abstract

Farm animals may harbor viral pathogens, some with zoonotic potential which can possibly cause severe clinical outcomes in animals and humans. Documenting the viral content of dust may provide information on the potential sources and movement of viruses. Here, we describe a dust sequencing strategy that provides detailed viral sequence characterization from farm dust samples and use this method to document the virus communities from chicken farm dust samples and paired feces collected from the same broiler farms in the Netherlands. From the sequencing data, *Parvoviridae* and *Picornaviridae* were the most frequently found virus families, detected in 85-100% of all fecal and dust samples with a large genomic diversity identified from the *Picornaviridae*. Sequences from the *Caliciviridae* and *Astroviridae* familes were also obtained. This study provides a unique characterization of virus communities in farmed chickens and paired farm dust samples and our sequencing methodology enabled the recovery of viral genome sequences from farm dust, providing important tracking details for virus movement between livestock animals and their farm environment. This study serves as a proof of concept supporting dust sampling to be used in viral metagenomic surveillance.

## 1. INTRODUCTION

Many emerging infectious diseases are of zoonotic origins ^1^. Approximately 70 % of zoonoses are proposed to originate from wildlife and outbreaks of MERS-CoV ^2^ from camels, Nipah virus ^3^ from bats and SARS-CoV-2 from minks ^4^ have shown that livestock and farmed animals can act as intermediate hosts and/or reservoirs which can sustain zoonotic transmission. In addition to passing viruses from wildlife to humans, livestock and farmed animals are also reservoir of zoonotic pathogens, such as bacteria *Coxiella burnetii (C. burnettii)* that causes Q-fever in goats, sheep and cattle ^5^, and avian influenza ^6,7^. Additionally, epidemiological studies have found evidence for an increased risk of respiratory disease in persons exposed to large-scale animal farming ^8,9^. In part, this can be attributed to farm dust exposure for which adverse health effects have been demonstrated ^10–12^, and the potential of zoonotic infections in these cases could be important yet under-reported. For instance, a Q-fever outbreak affecting approximately 4,000 people was reported in the Netherlands in 2007-2010 that was linked to circulation of *C. burnettii* in local dairy goat farms and dairy sheep farms ^13^. Q-fever was not notifiable in small ruminant farms in the Netherlands before the outbreak, making early detection of zoonotic transmission of *C. burnettii* challenging ^14^. Similarly, MERS-CoV had been circulating among dromedary camels long before it was recognized as a cause of severe respiratory disease in humans ^15^. These livestock-associated zoonotic disease outbreaks emphasize that continued surveillance of livestock is essential, especially when the intensification of livestock production is widely practiced to meet increasing food demands of growing populations ^16^.

Advances in next-generation sequencing (NGS) technology can provide a detailed description of the viral content of environmental samples and offers great potential for viral surveillance. Viral metagenomic approaches (using random priming rather than target specific primers) allow the detection of viruses present in a sample without prior knowledge of its presence ^17,18^ as well as characterization of complex viral communities ^19,20^. Indeed, reports of virus diversity in animals including bats ^21–23^, pigs ^24–26^, wild birds ^27,28^, and chickens ^29^ have been published since the development of NGS methods.

Chickens (*Gallus gallus domesticus*) are the most abundant domestic livestock in the world. According to the Food and Agriculture Organization of the United Nations (FAO), the global chicken population was >22 billion birds in 2017 ^30^. Chicken farming ranges from backyard to large-scale commercial farms that house ten-thousands of chickens. The Netherlands is the second largest agricultural exporter in the world ^31,32^. There are approximately 0.1 billion housed chickens and around 1,700 chicken farms in the Netherlands in 2020 ^33^. Chicken can carry various pathogens such as *Salmonella* ^*34*^, *Campylobacter* ^*35*^ and avian influenza viruses including subtypes that can cause severe infection in humans like H5N1 ^36^ and H7N9 ^6^; and it has been found that avian influenza virus can be transmitted to humans as a result of close exposure to contaminated chicken ^7^. Monitoring virus diversity in chickens and other livestock may be a key component for identifying potential zoonotic threats. Meanwhile, little is known about the virome composition of farm dust, although it has been suggested that farm dust might play a role in infectious disease transmission ^37,38^. Farm dust typically consists of fragments of animal feed, bedding material, animal feces, animal dander, mites and microorganisms ^39^, as well as fragments of poultry feathers in the case of poultry farm dust. The complex composition of dust particles allows them to accumulate and serve as a vector of biologically active material including antibiotic resistance genes, endotoxin, and, as shown here, complete viral genomes. Farm dust exposure may cause adverse health effects ^40,41^. Most studies on farm dust have focused on measuring antimicrobial gene sequences ^42^ and endotoxins ^10,12^, with no published reports documenting the viral sequence content of farm dust samples.

Here, we describe a metagenomic NGS characterization of virus diversity in a paired sample set of farm dust and chicken feces collected at multiple time points in three commercial chicken farms across the Netherlands. We hypothesized that the viral sequences identified from dust samples would share similarities with viral sequences detected from chicken fecal samples from the same farms. We develop and present a sequencing methodology that enabled recovery of virus genome sequences from farm dust samples. We show that the chicken fecal virome and farm dust virome show similar patterns, demonstrating the potential of using dust as a part of surveillance matrix for monitoring farm virus. This study provides useful insights in understanding the virus communities in chicken and surrounding farm environments and provides an important tool for monitoring viral movement by dust, a previously underappreciated source of viral movement through our environment.

## 2. RESULTS

### 2.1 Comparative virus detection in chicken farm dust samples and paired feces

A total of 46 individual farm dust samples and 56 pooled chicken feces were collected. Five chicken fecal samples were excluded due to insufficient cDNA concentration for library preparation. Individual farm dust samples collected at the same sampling moment from the same farm were pooled together during sample processing for sequencing, making up a final set of 13 pooled farm dust samples. Metagenomic deep sequencing of pooled chicken farm dust samples (N=13) and pooled chicken feces (N=51) generated ca 1-3 million paired-end short reads per sample. The negative blank control did not yield sufficient nucleic acid to be analyzed, indicating that consumables and reagents used were free of detectable viral materials.

The distribution of viral contigs from different virus families detected in chicken feces and chicken farm dust samples were generally similar **(Figure 1**). The number of contigs detected in *Picornaviridae* (farm dust samples: 199; chicken feces; 914) was the highest in both chicken farm dust and chicken feces samples, followed by *Parvoviridae* (farm dust samples: 183; chicken feces: 293), *Astroviridae* (farm dust samples: 135; chicken feces: 267) and *Caliciviridae* (farm dust samples: 85; chicken feces: 226). Interestingly, more DNA virus contigs were generally detected in chicken farm dust samples when comparing to that of chicken fecal samples, including adenoviruses, genomoviruses, smacoviruses and circoviruses.

**Figure 1.**
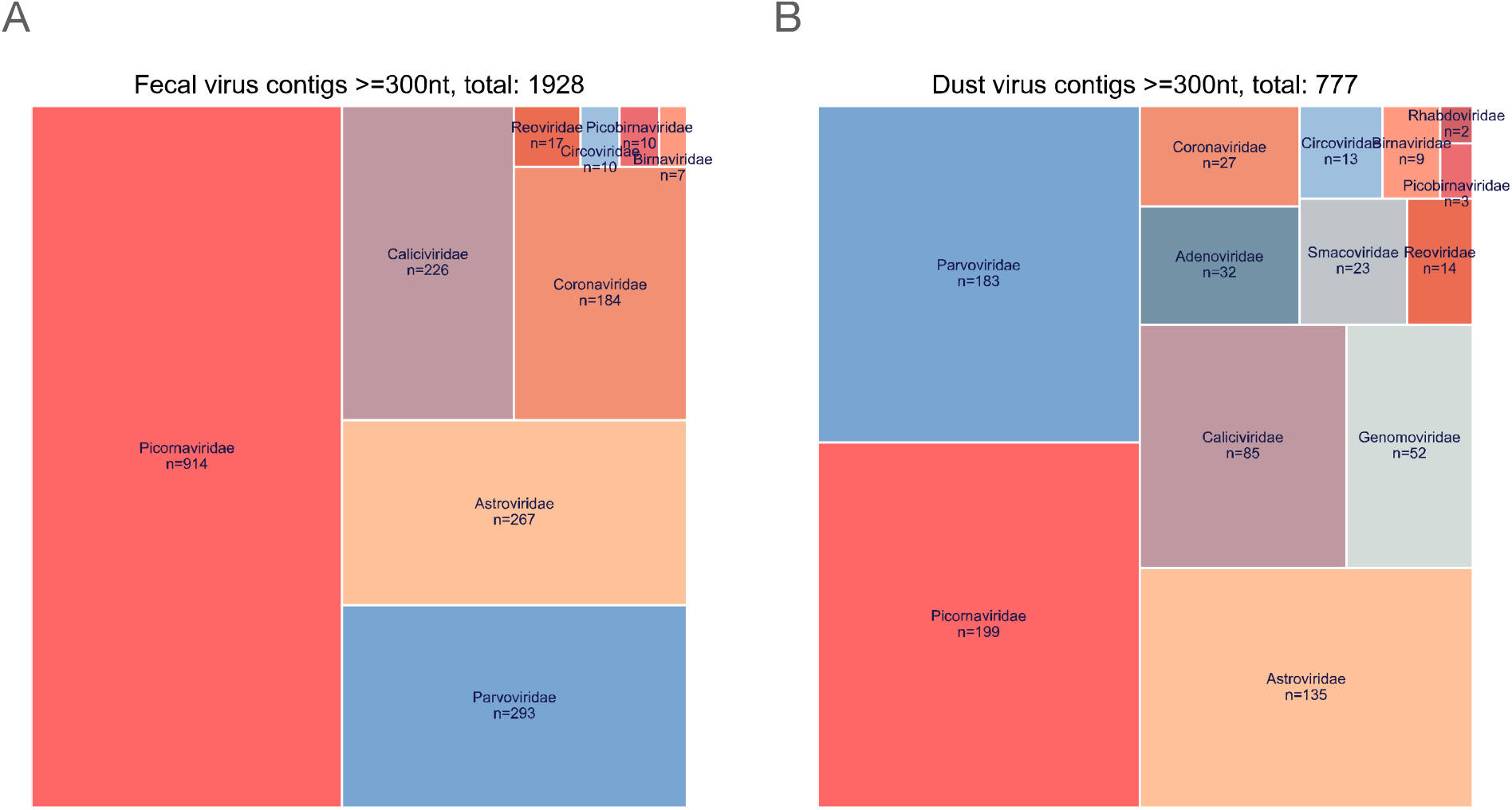
An overview treemap of the number of viral contigs detected in all chicken fecal samples (panel A) and all chicken farm dust samples (panel B). The size of each virus family sector is proportional to the number of contigs detected in the family (=number of contigs in this family/total number contigs observed). Only viral contigs with minimum length of 300 nt, at least 70 % identity to the reference sequences and a detection e-value threshold of <1×10^−10^ when comparing with the closest reference sequence in our database were included

From the total of 64 chicken farm dust samples (prefix: V_E; N=13) and chicken feces (prefix: V_M; N=51) analyzed, viral sequences belonging to the family *Parvoviridae* were consistently detected in all samples, followed by *Picornaviridae* with a 85% and 100% detection rate in chicken dust samples and chicken feces respectively **(Table 1)**. *Astroviridae* were the third most commonly detected virus family detected in 85% of farm dust samples and 82% chicken feces, followed by *Caliciviridae* which were present in 69% (farm dust samples) and 78% (chicken feces) detection rate. The overall pattern of virus detection observed in dust samples was similar to that in chicken feces, some more variability was observed within farms over time (**Table 1 and Figure S1**).

**Table 1.** Detection of viral sequences from different eukaryotic virus families in chicken feces and farm dust samples

| Sample types |  | Chicken feces (N=51) | Farm dust (N=13) |
| --- | --- | --- | --- |
| Virus family | *Astroviridae* | 42 (82%) | 11 (85%) |
|  | *Caliciviridae* | 40 (78%) | 9 (69%) |
|  | *Coronaviridae* | 19 (37%) | 3 (23%) |
|  | *Parvoviridae* | 51 (100%) | 13 (100%) |
|  | *Picobirnaviridae* | 8 (16%) | 2 (15%) |
|  | *Picornaviridae* | 51 (100%) | 11 (85%) |
|  | *Reoviridae* | 7 (14%) | 3 (23%) |

N is the total number of the samples. The value represents the number of viral sequences with minimum amino acid identity of 70%, minimum length of 300 nt and an e-value threshold of $1 \times 10^{-10}$ were included. The percentage is the detection percentage of the virus family in our samples.

Although the number of *Parvoviridae* positive samples was slightly higher than for *Picornaviridae*, the actual number of *Picornaviridae* sequence contigs detected was the highest (**Figure 1 and 2**). Within the *Picornaviridae* family, *Sicinivirus* sequences were consistently detected in 69% of farm dust samples and in all chicken feces (**Table 2**), followed by *Anativirus* (farm dust samples: 61%; chicken feces: 70%), and *Megrivirus* (farm dust samples: 46%; chicken feces: 43%). Of note, the number of contigs detected in different picornavirus genera corresponded to the detection pattern (**Figure 3 & Figure S2**). In particular, *Anatvirus, Gallivirus, Megrivirus* and *Sicinivirus* sequences were detected in all three farms, while *Orivirus* was only identified in farm F_01 and F_03 and *Avisivirus* was only found in farm F_03 (**Figure S2**). When looking at the virus family level, we did not observe any substantial differences in overall viral contents (**Figure 2**) and virus-specific detection (**Figure S1**) of samples from different farms and from different production cycles (different chicken ages) within farms.

**Table 2.** Detection of viral sequences from picornavirus genera in chicken feces and farm dust samples

| Sample types |  | Chicken feces (N=51) | Farm dust (N=13) |
| --- | --- | --- | --- |
| Picornavirus genera | *Anativirus* | 36 (70%) | 8 (61%) |
|  | *Avisivirus* | 3 (5%) | 0 (0%) |
|  | *Gallivirus* | 16 (31%) | 4 (30%) |
|  | *Megrivirus* | 22 (43%) | 6 (46%) |
|  | *Orivirus* | 8 (15%) | 0 (0%) |
|  | *Sicinivirus* | 51 (100%) | 9 (69%) |

N is the total number of the samples. The value represents the number of viral sequences with minimum amino acid identity of 70%, minimum length of 300 nt and an e-value threshold of $1 \times 10^{-10}$ were included. The percentage is the detection percentage of the picornavirus genera in our samples.

**Figure 2.**
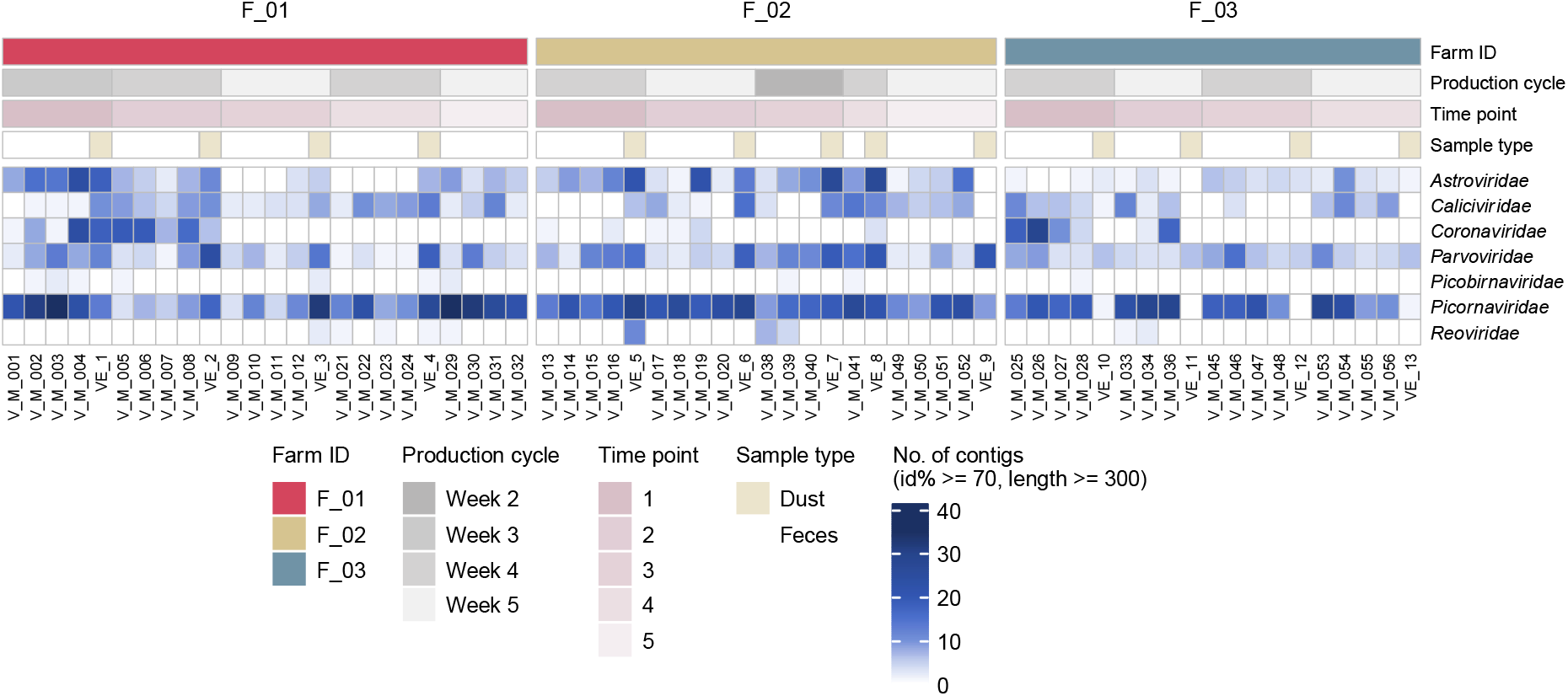
An overview of virus contigs detected in chicken feces and chicken farm dust samples. Each column represents one sample. Associated metadata is shown at the top panel. The color intensity of the heatmap (bottom panel) is determined by number of contigs with minimum length of 300 nt/bp, at least 70% identity and an e-value threshold of 1×10^−10^ when comparing with closest reference in our database. Sample order is assorted by farm ID, sampling time point. Time point is an arbitrary number. Samples were collected at the same time and place when both Farm ID and Time point match.

**Figure 3.**
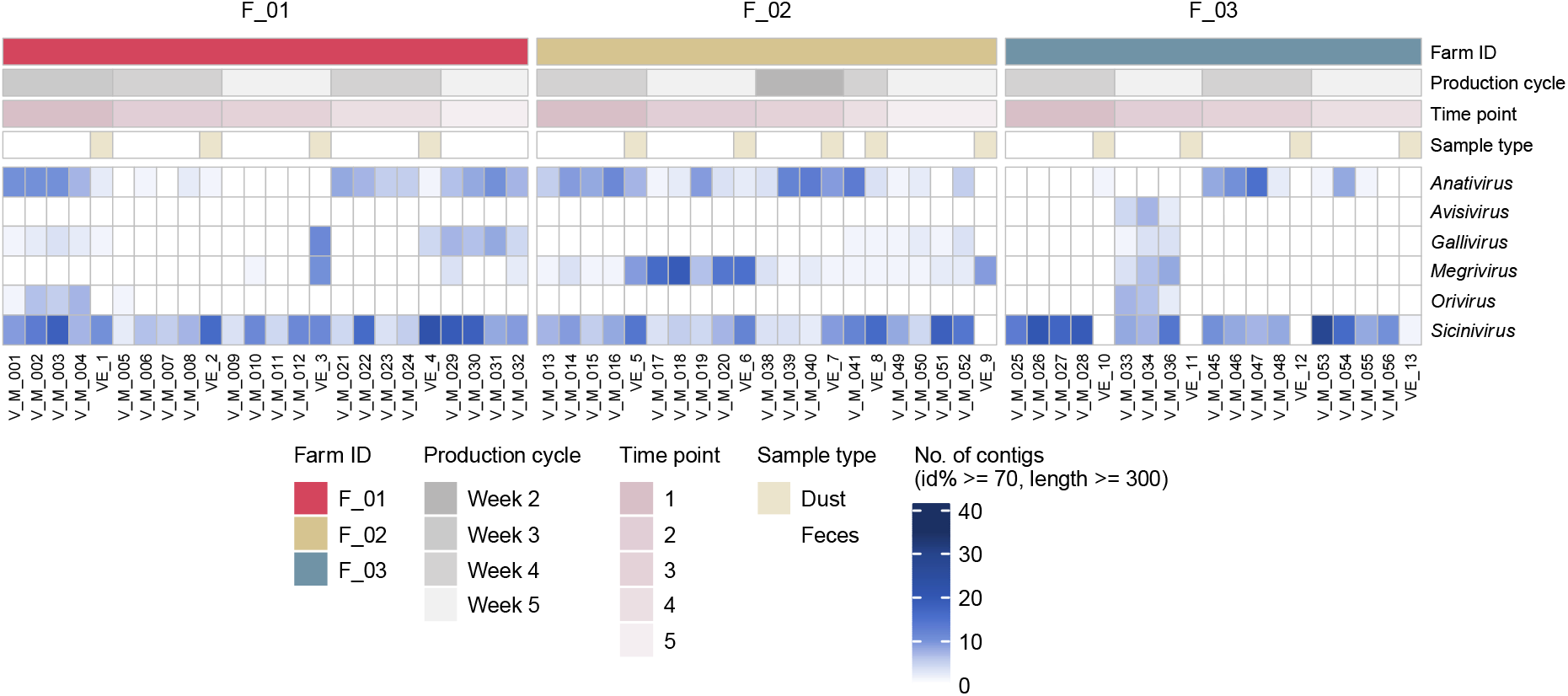
An overview of picornavirus contigs detected in chicken feces and chicken farm dust samples. Each column represents one sample. Associated metadata is shown at the top panel. The color intensity of the heatmap (bottom panel) is determined by number of contigs with minimum length of 300 nt, at least 70% identity and an e-value threshold of 1×10^−10^ when comparing with closest reference in our database. Sample order is assorted by farm ID, sampling time point. Time point is an arbitrary number. Samples were collected at the same time and place when both Farm ID and Time point match.

### 2.2 Genomic diversity in viral sequences identified from chicken farm dust samples and paired feces

#### 2.2.1 Picornaviridae

Phylogenetic construction was performed to compare the reported *Picornaviridae* sequences from 5 genera (*Avisvirus, Anativirus, Gallivirus, Megrivirus* and *Sicinivirus*) with global sequences. Forty-six picornavirus sequences with ≥ 80% genome coverage (farm dust samples: 8; chicken feces: 38) were included in the phylogenetic analysis of the polyprotein region. The maximum-likelihood (ML) phylogenetic tree (**Figure 4A**) showed that the sequences identified from farm F_03, albeit small number (N=12), belonged to all 5 different genera *Avisvirus, Anativirus, Gallivirus, Megrivirus* and *Sicinivirus*, suggesting a high viral diversity circulating among chicken in this farm. Interestingly, farm F_02 with the highest number of picornavirus contigs identified (N=31) has a moderate level of viral diversity, with these contigs belonging to 3 genera *Anativirus, Sicinivirus* and *Megrivirus*. Only 3 picornavirus genomic sequences identified were from farm F_01, belonging to *Sicinivirus* (N=1) and *Anativirus* (N=2). *Sicinivirus* and *Anativirus* are the most commonly detected virus genera found with virus sequences detected in all 3 farms, while sequences of *Gallivirus* and *Avisivirus* were less commonly detected, found in only 1 farm, farm F_03.

**Figure 4.**
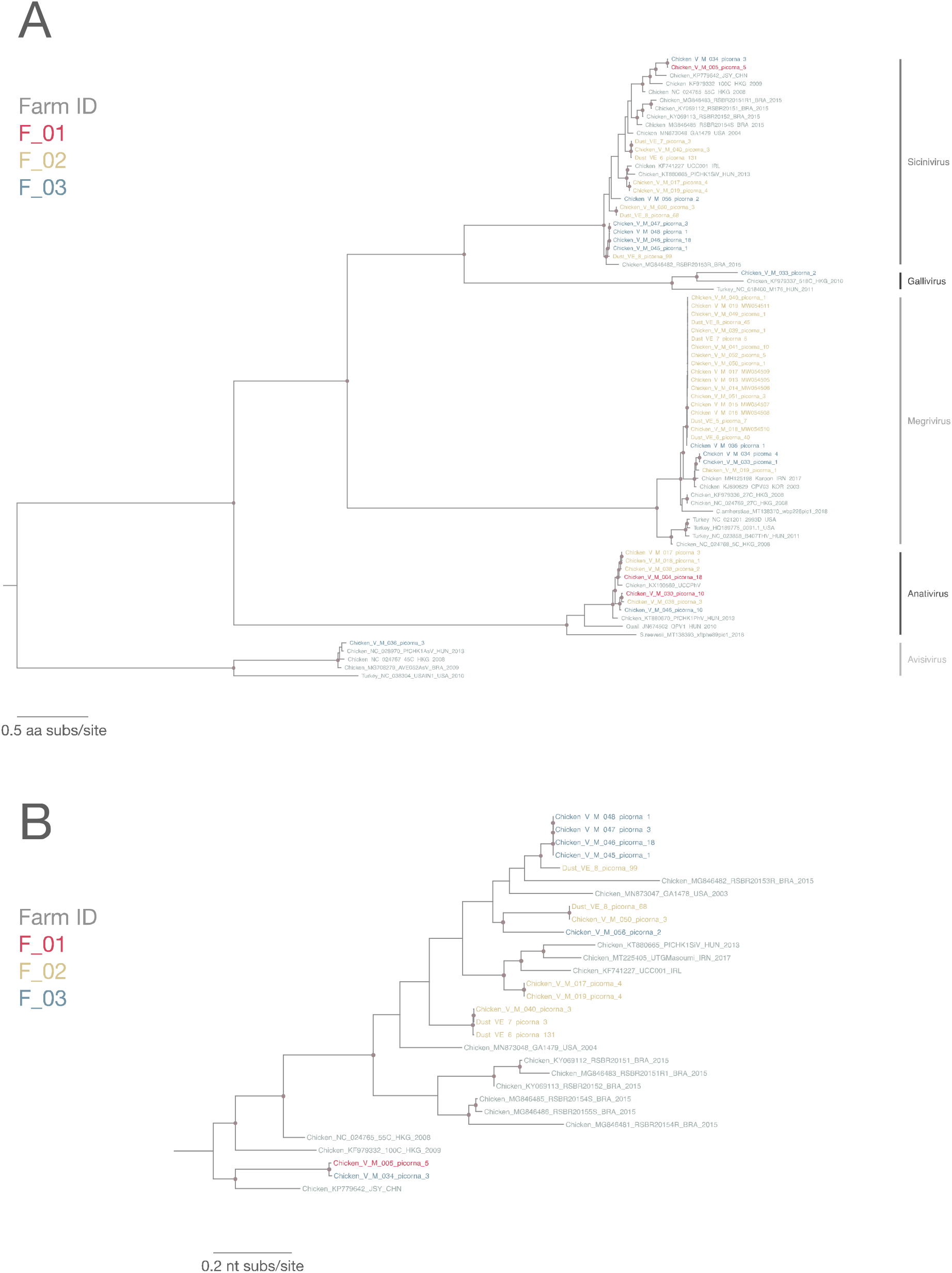
A. Maximum-likelihood phylogenetic tree of 76 (sequences from this study: 46; reference sequences: 30) picornavirus partial polyprotein amino acid sequences (3786 sites) reconstructed using a best-fit evolutionary model LG+FC+R10. For clarity, only bootstrap supports >= 70 are shown as a navy-blue node point. Tip labels of our samples are color-coded by corresponding farm ID. Reference sequences are in grey. Tree was mid-point rooted for clarity. B. Maximum-likelihood phylogenetic tree of 30 *Sicinivirus* partial polyprotein nucleotide sequences (sequences from this study: 15; reference sequences: 15; 8672 sites) reconstructed using a best-fit evolutionary model GTR+F+R10. For clarity, only bootstrap supports >= 70 are shown as a navy-blue node point. Tip labels of our samples are color-coded by corresponding farm ID. Reference sequences are in grey and tree was mid-point rooted for clarity.

Genome sequences detected from farm dust samples belonged to *Sicinivirus* and *Megrivirus* from farm F_02 that clustered with sequences identified in chicken feces from the same farm (**Figure 4A**). Viral sequences in the *Sicinivirus* genus were detected in 69% of dust samples and all chicken feces samples with the highest number of viral contigs identified in each sample as compared to other genera in the *Picornaviridae* family. Slightly longer branches in the *Sicinivirus* sequences in the ML tree indicates higher diversity within this *Sicinivirus* clade (**Figure 4A**). We further reconstructed a ML tree for all identified *Sicinivirus* polyprotein nucleotide sequences (N=15; farm dust samples: 4; chicken feces: 11) in this study and compared them with global reference sequences (**Figure 4B**). Two sequences from chicken feces (V_M_005_picorna_5 from farm F_01 and V_M_034_picorna_3 from farm F_03) were most distinct from the rest of the identified sequences (sharing only 69.4-72.3% nt identity) and were most closely related to strain JSY previously identified in China (78.9-79.0% nt identity). Other *Sicinivirus* sequences from the same farm often formed a monophyletic lineage, suggesting within-farm *Sicinivirus* sequences are highly genetically similar. Viral sequences identified from chicken feces and farm dust samples consistently clustered with each other forming their own sub-clusters within major clades in the ML trees, indicating high similarity between sequences from feces and dusts and that they are more closely related as compared to global reference sequences.

#### 2.2.2 Astroviridae

Sixteen astrovirus sequences with ≥ 80% genome coverage were identified in this study, among which only 1 astrovirus sequence was found in farm dust (VE_7_astro_14; farm F_02). There was only 1 astrovirus genomic sequence from farm F_01. RNA-dependent RNA polymerase (RdRp) and capsid regions of these sequences were extracted for phylogenetic analyses (**Figure 5A and 5B**, respectively). ML trees of both regions showed that these 16 astrovirus sequences belonged to two distinct lineages, A1 and A2 (**Figure 5)**. Sequences from samples from farm F_02 and F_03 were found in both lineages A1 and A2, suggesting co-circulation of different astrovirus strains in both farms. The 12 sequences from lineage A1 were most closely related to avian nephritis virus strains previously reported in China and Brazil (GenBank accession number: <u>HM029238</u>, <u>MN732558</u> and <u>MH028405</u>), while the 4 sequences from lineage A2 were more closely related to each other and clustered with chicken astroviruses previously identified in Malaysia, Canada and the USA (GenBank accession number: <u>MT491731</u>-<u>2</u>, <u>MT789782</u>, <u>MT789784</u> and <u>JF414802</u>). The 12 sequences from lineage A1 shared 96-100% amino acid (aa) identity in the RdRp region while 4 sequences from lineage A2 shared a higher 99-100% aa identity. For the less conserved capsid region, three small sub-lineages can be observed within lineage A1 (**Figure 5B**). Notably, the only astrovirus sequence found in dust (strain VE_7_astro_14) was identical to astrovirus sequence recovered from its paired chicken feces sample (strain V_M_038_astro_4) at the same farm.

**Figure 5.**
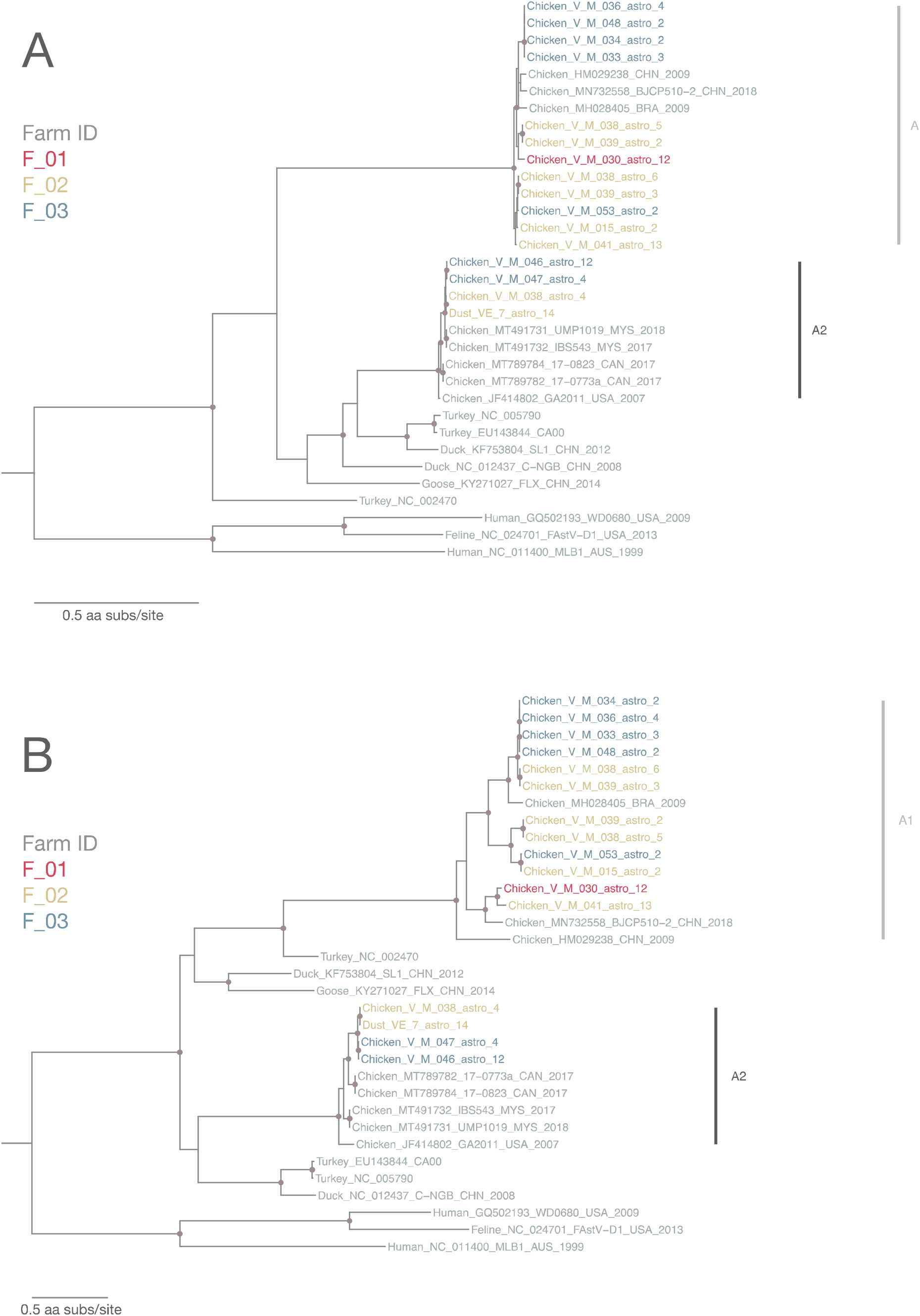
A. Maximum-likelihood phylogenetic tree of 33 astrovirus partial RdRp amino acid sequences (sequences from this study: 16; reference sequences: 17; 548 sites) reconstructed using a best-fit evolutionary model LG+FC+I+G4m. B. Maximum-likelihood phylogenetic tree of 33 astrovirus partial capsid amino acid sequences (sequences from this study: 16; reference sequences: 17; 908 sites) reconstructed using a best-fit evolutionary model LG+FC+I+G4m. For clarity, only bootstrap supports >= 70 are shown as a navy-blue node point. Tip labels of our samples are color-coded by corresponding farm ID. Reference sequences are in grey and tree was mid-point rooted for clarity.

#### 2.2.3 Caliciviridae

Similar to astrovirus distribution in feces versus dust, 13 out of 14 calicivirus sequences identified (with ≥ 80% genome coverage) were recovered from fecal samples with only 1 calicivirus sequence identified from a dust sample. Partial ORF1 polyprotein region of these sequences was extracted to construct ML tree for comparing the local calicivirus with global sequences. The ML tree showed that the reported calicivirus sequences were from two distinct lineages (C1 and C2) (**Figure 6**). All calicivirus sequences from farm F_02 and F_03 belonged to lineage C1, while sequences from F_01 belonged to both lineages C1 and C2. The 11 sequences in lineage C1 shared 98.7-100% aa identity despite being collected from different farms and they shared 97.2-99.1% aa similarity with two partial ORF1 sequences identified in Germany in 2010 (GenBank accession number: <u>JQ347523</u> and <u>JQ347527</u>). BLAST searches and multiple sequence alignment indicated that these 11 local calicivirus sequences are the first report of complete ORF1 sequence of that particular chicken calicivirus lineage. The only calicivirus sequence from dust (VE_8_calici_66) clustered in lineage C1. The 3 sequences in lineage C2 are highly similar, sharing 99.9-100% aa identity and were all collected from farm F_01 at two time points. These 3 sequences shared 93.8-96.8% aa similarity with reference sequences from Germany, USA, Korea and Brazil (*Bavaria virus, Bavovirus* genus, GenBank accession number: <u>HQ010042</u>, <u>MN810875</u>, <u>KM254170</u>-<u>1</u>, <u>MG846433</u>-<u>4</u>).

**Figure 6.**
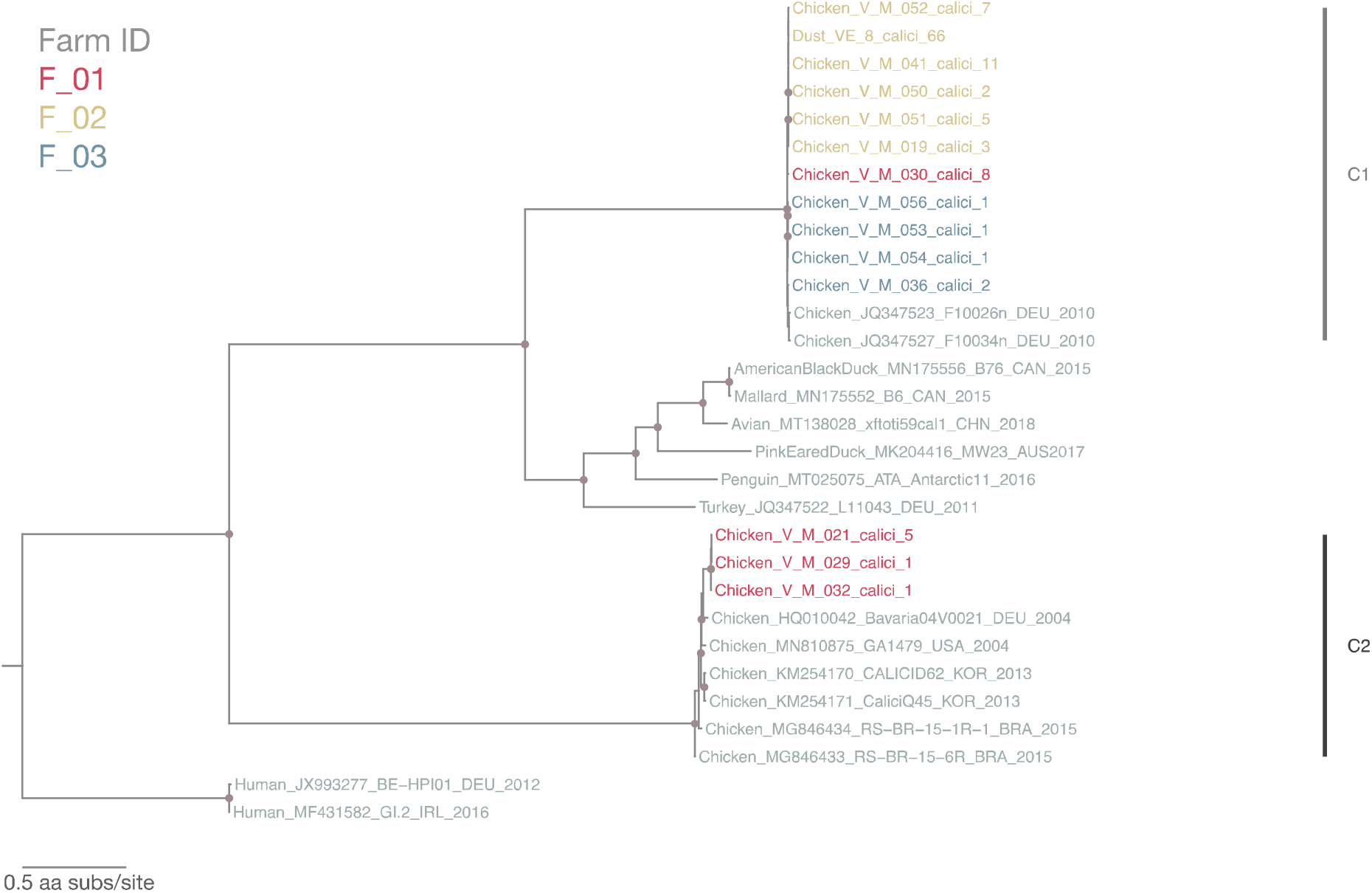
Maximum-likelihood phylogenetic tree of 30 calicivirus partial polyprotein amino acid sequences (sequences from this study: 14; reference sequences: 16; 2794 sites) reconstructed u best-fit evolutionary model LG+FC+R10. For clarity, only bootstrap supports >= 70 are shown as blue node point. Tip labels of our samples are color-coded by corresponding farm ID. Reference sequences are in grey and tree was mid-point rooted for clarity.

#### 2.2.4 Parvoviridae

A total of 34 parvovirus sequences with ≥ 80% genome coverage were identified (chicken farm dust: 8; chicken feces: 28). The amino acid sequences of the nonstructural protein (NS1) region were extracted for ML tree reconstruction. Two distinct lineages (P1 and P2) were observed in the ML tree supported by >70% bootstrap value (**Figure 7**). All but one sequences from farm F_01 and F_03 were clustered in lineage P1 and lineage P2 respectively, while sequences from farm F_02 belonged to both lineages. The 16 sequences in lineage P1 shared 99.5%-100% aa identity and they shared 97.3-99.6% aa identity with sequences from Switzerland and Brazil (GenBank accession number: <u>OM469025</u>, <u>OM469088</u>, <u>OM469107</u> and <u>MG846442</u>-<u>3</u>). The 18 sequences in lineage P2 shared 99.4-100% aa identity with each other and 98.7-99.9% aa identity to sequences from Switzerland, Brazil, Korea and Canada (GenBank accession number: <u>OM469032</u>, <u>OM469027</u>, <u>MG846441</u>, <u>KM254174</u> and <u>MW306779</u>). The inter-lineage aa identity between lineage P1 and P2 ranged from 73.7-74.4%.

**Figure 7.**
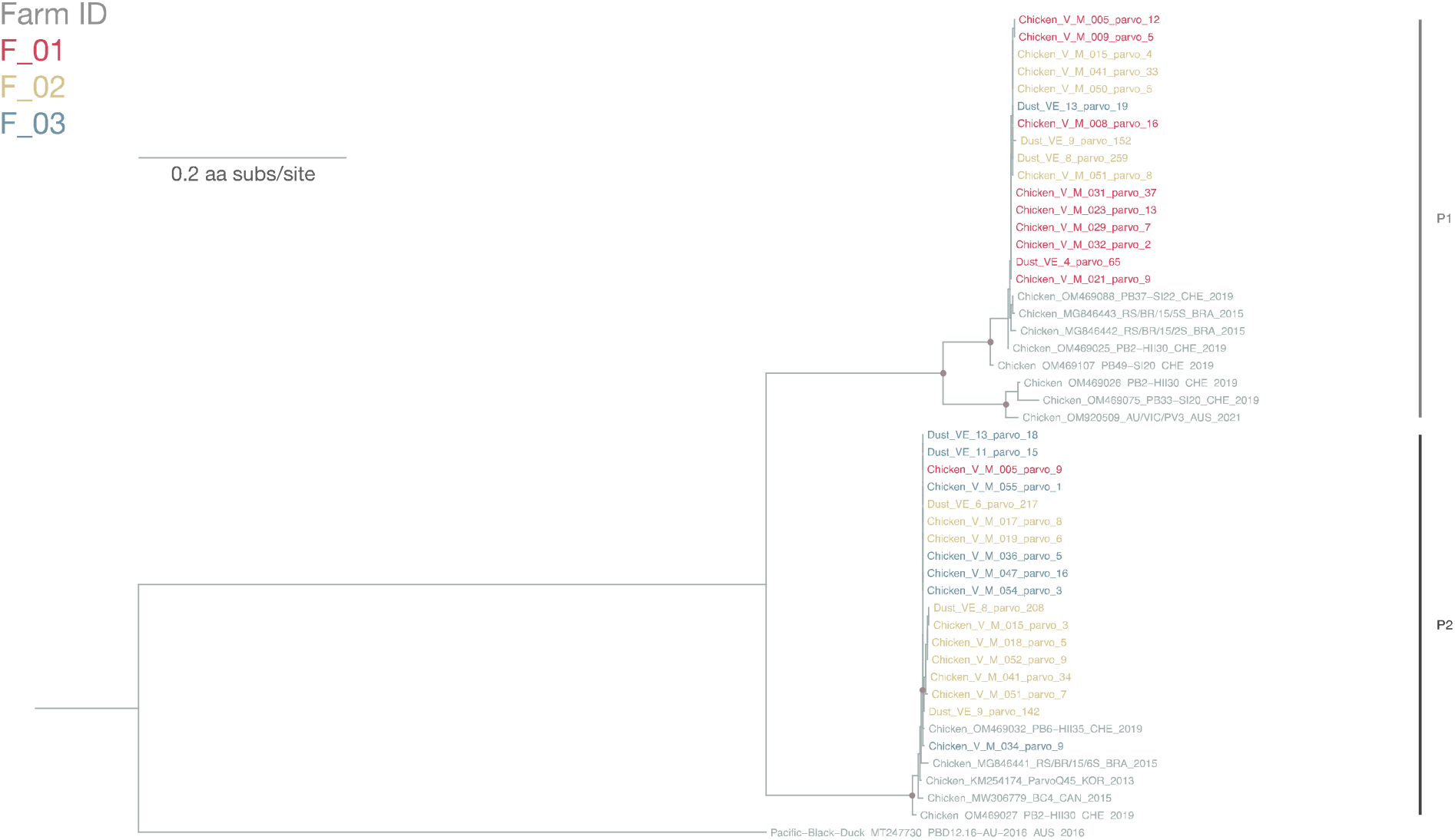
Maximum-likelihood phylogenetic tree of 48 parovirus partial NS1 amino sequences (sequences from this study: 34; reference sequences: 14; 700 sites) reconstructed using a best-fit evolutionary model JTT+FC+R2. For clarity, only bootstrap supports >= 70 are shown as a navy-blue node point. Tip labels of our samples are color-coded by corresponding farm ID. Reference sequences are in grey. The tree is rooted to a chaphamaparvovirus 1 strain, named PBD12.16-AU-2016, identified from Pacific black duck (GenBank accession number: MT247730).

#### 2.2.5 Other virus families

Three nearly-complete coronavirus genome sequences were identified in 3 chicken feces (V_M_001, V_M_002 and V_M_003) from farm F_01. BLAST analyses showed that these sequences are infectious bronchitis virus (IBV) strains and share 99.92-99.95% identity at the nucleotide level with IBV vaccine strain 4/91 (GenBank accession number: <u>KF377577</u>), suggesting these IBV may possibly be viral shedding following vaccination or environmental contamination as vaccines are generally provided in drinking water.

### 2.3 Mash analysis

The phylogenetic comparison of large contig sequences presented above support the findings about the similarities between sequences from dust samples and possible farm animal sources. However, additional information can be obtained with global analyses that would compare all available sequence data from the samples. A method that allows a comparison of the large amount of unclassified sequences that are commonly observed in NGS data would provide additional information. We therefore used a kmer comparison tool <u>Mash</u> ^43,44^ that prepares a hash description of all kmers of a given length and then allows rapid quantitative comparison of similarity of the kmers sets generated from different sequenced samples. We used the Mash distance function to calculate a Jaccard distance value for all pairs of dust sequence samples.

We compared all assembled viral contigs of the dust samples. Figure 8A shows the Jaccard distance between all 13 dust samples from 3 farms, colored by farm source. In all comparisons, pairs of dust samples from the same farm have lower Jaccard distance values (= greater similarity, darker blue in the figure) than inter-farm samples. It is also apparent that farm F_01 and farm F_02 were slightly more related to each other than either were to farm F_03. Similar patterns were obtained using all quality-controlled short read data from each dust sample (**Figure 8B**).

**Figure 8.**
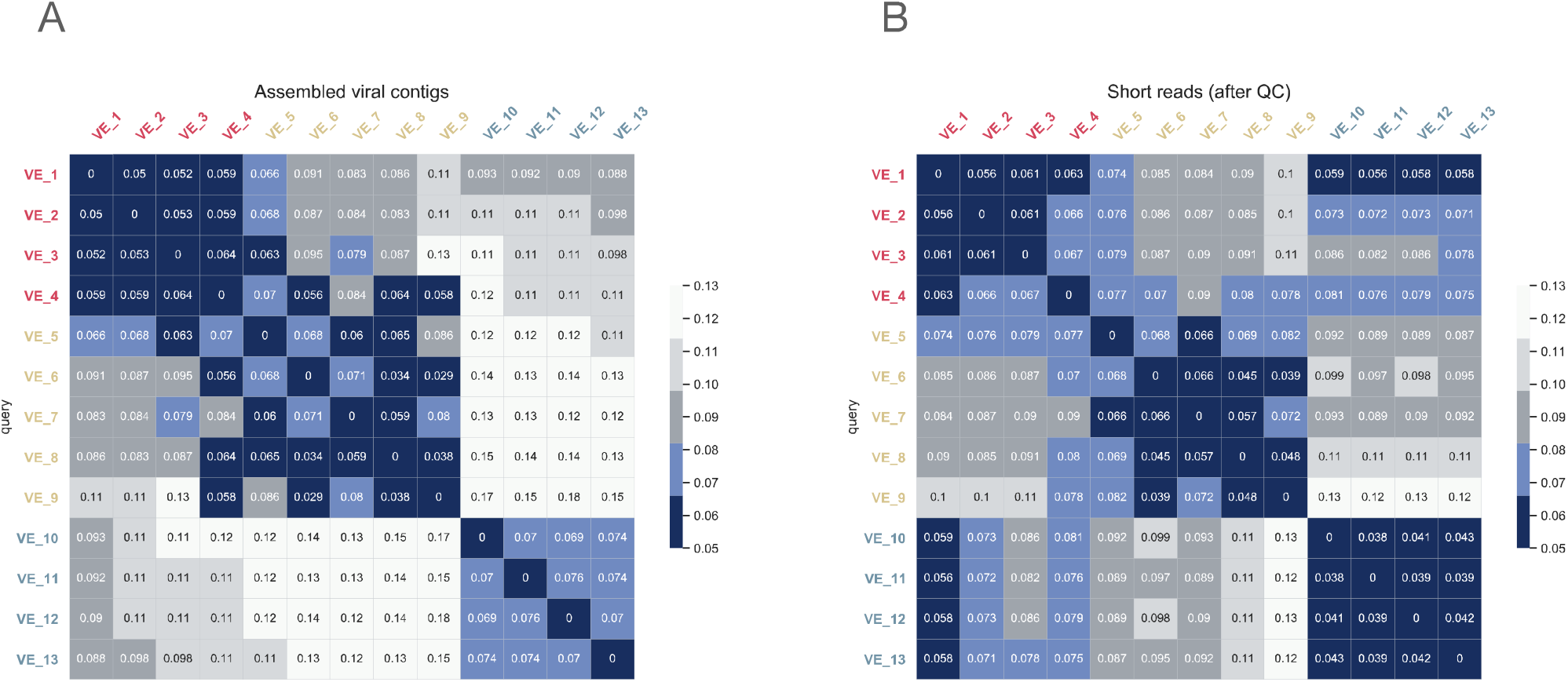
A. Jaccard distance analysis of assembled viral contigs of 13 pooled farm dust samples. B. Jaccard distance analysis of quality-controlled short reads of 13 pooled farm dust samples. The color intensity of the heatmaps is determined by MASH distance. Corresponding MASH distances are also shown in the heatmap. Sample labels are color-coded by farm ID and farm ID was added as prefix for clarity.

## 3. DISCUSSION

Animals including livestock kept in close proximity to humans can act as intermediate hosts and/or sources for zoonotic disease transmission ^45^. The current COVID-19 pandemic has raised awareness of zoonotic risks and has prompted the need for more proactive surveillance strategies involving animal surveillance, which could potentially guide zoonotic outbreak preparedness. Chickens are the most abundant livestock in the world ^30^ and have been identified as reservoirs for various zoonotic pathogens ^35,36^. In this study, we employed viral metagenomic deep sequencing to document the viruses present in farm dust samples and paired chicken feces collected within the same farms. These data revealed potential virus flow between chickens and their farm environment, highlighting the important role of dust as part of surveillance matrix to bring useful insights into the equation for monitoring virus flows at the animal-environment interface.

In this study, we described a random-primed viral metagenomic deep sequencing strategy to obtain viral sequences from dust samples. We were able to obtain long viral sequences providing ≥ 80% viral genome coverage from farm dust samples. Farm dust is known to be associated with adverse respiratory effects observed in farm workers with prolonged exposure ^40^; however, whether viruses play a role is thus far unknown due to a knowledge gap in virus detection and virus diversity in farm dust. Characterizing farm dust viromes will aid in investigating possible health effects of occupational and environmental exposure to virus-containing farm dust. The method we have developed provides a platform for future surveillance in farm animals and environmental samples.

Viral contig distribution of chicken dust samples was in general comparable to that of chicken fecal samples. Viral sequences from *Picornaviridae, Parvoviridae, Astroviridae* and *Caliciviridae* were the most commonly detected in both sample types (**Figure 1**), although a higher abundance of DNA virus sequences was interestingly observed in dust samples. Our phylogenetic analyses indicated that for all three farms monitored, viruses identified from farm dust samples were genetically closer to viruses identified from chicken feces collected from the same farm than from the other two farms in the study. Furthermore, the Jaccard similarity analysis (**Figure 8**) further showed that for both known and unknown sequences the dust sequences were closer to other samples from the same farm, indicating some source specificity in the sampling method. This observation is in agreement with a previous study that reported the correlation found in the bacterial antimicrobial resistomes between animal feces and farm dust ^46^. Although infectivity of the viruses identified was not determined in our study, the presence of nearly complete viral genome sequences and the complex protein/lipid/nucleic acid composition of dust makes it likely that dust indeed contains infectious virus particles. Future investigation on the infectivity of viruses in farm dust will provide useful insights in mapping virus flows between chickens and their surrounding environment, which collectively would guide future outbreak control strategies within and between farms.

Viral sequences identified in chicken feces from different farms were phylogenetically closely related to each other and formed sub-lineages or monophyletic lineages when comparing to global related sequences. This could possibly be explained by the pyramid structure of the broiler production system, as chicks found at each farm were all bought from a few suppliers; in other words, chicks could have been infected from the supplier and then brought the virus to different broiler farms with further spread within the farms, leading to similar viruses circulating in different broiler farms. Of note, the pyramid production structure was also suggested as a possible transmission route for ESBL/pAmpC producing bacteria between broiler farms ^47^. Future characterization of virus diversity of chickens from the top part of the pyramid (suppliers) is needed to confirm the speculation.

Viral sequences belonging to five genera of the *Picornaviridae* family were identified in our study, suggesting a great diversity of picornaviruses circulating in chickens, which is consistent with previous studies that detected various picornavirus genera in chicken respiratory samples ^48,49^. Astroviruses identified in this study clustered in two distinct phylogenetic lineages. All chicken astroviruses are grouped to one single species according to the current taxonomy assignment from International Committee on Taxonomy of Viruses ^50^. Current classification of avastrovirus is based on phylogenetic analysis of capsid (ORF2) amino acid sequences ^51^. More comprehensive assignment of genotypes is hindered by the limited number of complete capsid sequences. A similar observation was seen for *Caliciviridae*, where we identified two types of caliciviruses forming two distinct lineages in the ML tree. Lineage C2 is closely related to the *Bavovirus* genus, while the sequences from C1 are most related to an unclassified chicken calicivirus strain F10026n ^52^ that only possessed a partial ORF1 sequence. Our data have provided 16 astrovirus and 14 calicivirus sequences, which helps expand the current knowledge of chicken astrovirus and calicivirus diversity and could potentially contribute to improved taxonomic assignment.

We observed that viral sequences in the *Parvoviridae* and *Picornaviridae* families were consistently detected in chicken feces and farm dust samples at multiple time points in all three farms. These virus families could potentially be used as signature markers when monitoring chicken and farm dust exposure. Additionally, these viruses could also be used as markers to assess the efficiency of disinfection procedures in farms and their impact on surrounding environments although stabilities of different viruses may differ due to different structures and we caution that viral activity assays have not yet been performed on this material. Further and perhaps larger scale of longitudinal surveillance with the inclusion of samples from layer hens and more environmental samples from surrounding areas, as well as samples from other bird species are certainly necessary to confirm our observation and the applicability of using these identified viruses as signature virus fingerprint.

Our study has several limitations. Although we collected chicken feces and farm dust samples longitudinally, limited number of samples per each time point especially for farm dust samples has hindered in-depth statistical comparisons between samples collected from different time points. An important limitation that should be considered is that the fecal and dust samples were each generated from pooled material, and dust by its very nature is a pooled sample (i.e. possibly containing sequences from multiple individual infections). The possibility that the resulting assembled genomes from these fecal and dust samples may be chimeras assembled from multiple individual infections should be considered in interpreting the results. This means that although one might intuitively expect identical sequences identified from the dust originating from the same source as the fecal sample, the technology and the pooling means that identical genome sequences may be difficult to find even though the sources of the viruses could be the same. Virologically important viruses should be examined using alternate methods such as direct isolation from individually infected animals. Chicken fecal samples and dust samples were processed slightly differently because of different sample nature and reagent availability, which could potentially lead to minor differences in detection sensitivity. Our finding showed that viral sequence content found in dust and chicken feces are relatively similar, suggesting any potential bias is minimal.

In conclusion, we provide a dataset of viral sequences generated from farm dust samples and feces and show a similar pattern between viral sequences from dust and feces samples in terms of composition and genetic similarity. These results support the idea that dust sampling can provide an accurate description of the farm virome and could potentially be incorporated as part of viral metagenomic surveillance matrix. In the long term, understanding virus flow between animals and humans is vital for identifying potential zoonotic threats and zoonotic outbreak control and preparedness.

## 4. MATERIALS AND METHODS

### 4.1 Sample collection

Pooled chicken feces and farm dust samples were collected in three commercial broiler farms in the Netherlands at 4-5 time points from May to August 2019. The three commercial broiler farms are located in three different regions in the Netherlands (Western part: Noord/Zuid-Holland region; Northern part: Friesland/Groningen region; Eastern part: Gelderland region). Detailed sampling strategy and sample metadata is described in **Table 3**. Each pooled poultry fecal sample was manually picked from the floor and contains fresh fecal material from 3-4 chicks. The chicks were not kept in cages but could walk freely. Farm dust samples were collected using a passive air sampling approach using electrostatic dustfall collectors (EDCs) ^53,54^. Electrostatic cloths were sterilized through incubation at 200°C for 4 hours. Sterilized electrostatic cloths were then fixed to a pre-cleaned plastic frame. EDCs were exposed for 7 days at 1 meter above the floor with the electrostatic cloths facing up in broiler farms to enable sampling of settling airborne dust instead of resuspended dust from the floor. EDCs were contained in a sterile plastic bag before and after sampling. All samples were transported under cold chain management and stored at -20°C/-80°C before processing.

**Table 3.**
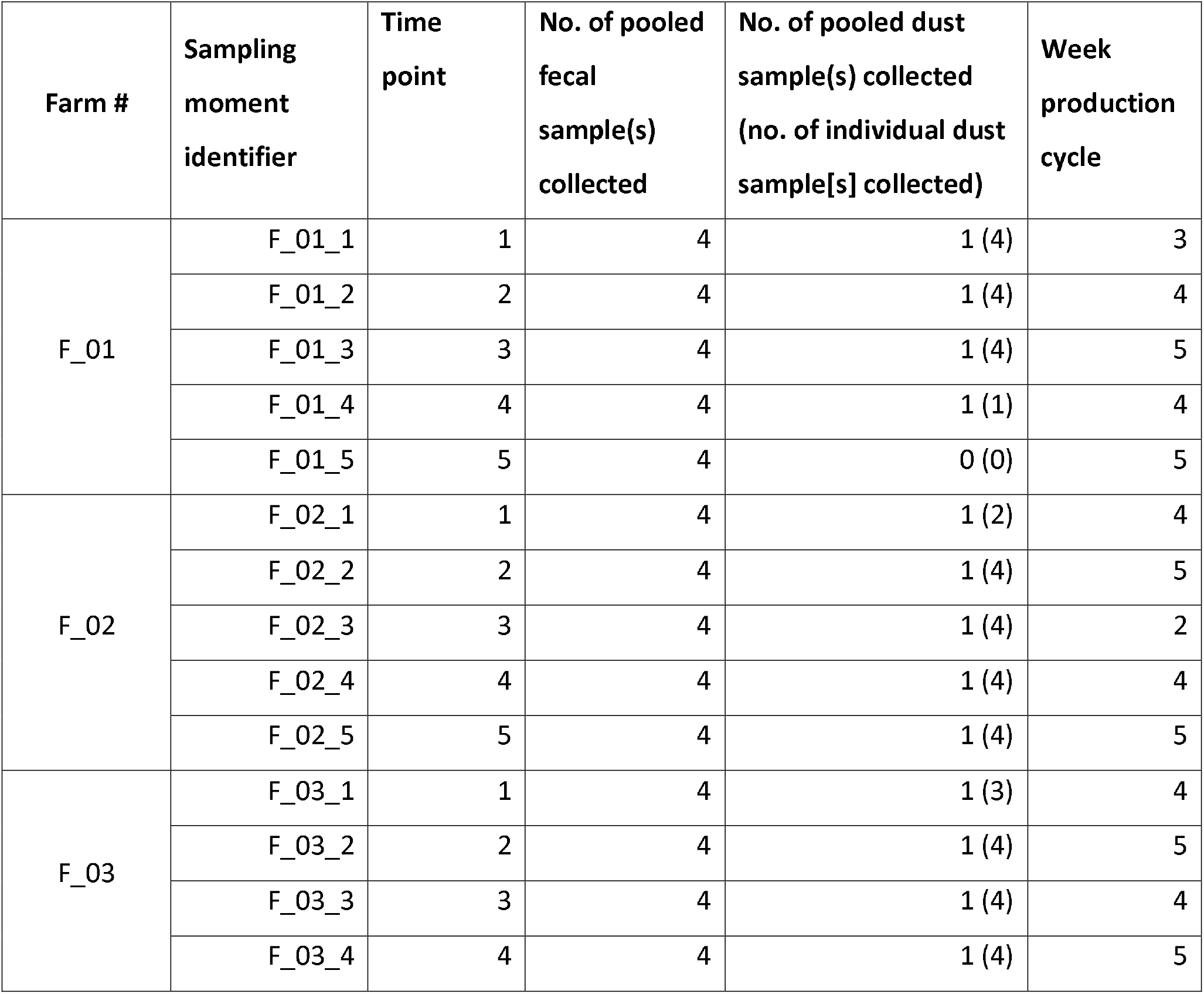
Detailed sampling strategy

| Farm # | Sampling moment identifier | Time point | No. of pooled fecal sample(s) collected | No. of pooled dust sample(s) collected (no. of individual dust sample[s] collected) | Week production cycle |
| --- | --- | --- | --- | --- | --- |
| F_01 | F_01_1 | 1 | 4 | 1 (4) | 3 |
|  | F_01_2 | 2 | 4 | 1 (4) | 4 |
|  | F_01_3 | 3 | 4 | 1 (4) | 5 |
|  | F_01_4 | 4 | 4 | 1 (1) | 4 |
|  | F_01_5 | 5 | 4 | 0 (0) | 5 |
| F_02 | F_02_1 | 1 | 4 | 1 (2) | 4 |
|  | F_02_2 | 2 | 4 | 1 (4) | 5 |
|  | F_02_3 | 3 | 4 | 1 (4) | 2 |
|  | F_02_4 | 4 | 4 | 1 (4) | 4 |
|  | F_02_5 | 5 | 4 | 1 (4) | 5 |
| F_03 | F_03_1 | 1 | 4 | 1 (3) | 4 |
|  | F_03_2 | 2 | 4 | 1 (4) | 5 |
|  | F_03_3 | 3 | 4 | 1 (4) | 4 |
|  | F_03_4 | 4 | 4 | 1 (4) | 5 |

### 4.2 Sample processing and metagenomic deep sequencing

Chicken fecal samples were processed as previously described ^55^. Briefly, chicken fecal suspension was prepared in Phosphate-buffered saline (PBS) and treated with TURBO DNAse (Thermo Fisher, USA) at 37 °C for 30 minutes, and then subjected to total nucleic acid extraction using QIAamp viral RNA mini kit (Qiagen, Germany) according to manufacturer’s instruction without addition of carrier RNA. For dust samples, electrostatic cloths were incubated in 3% beef extract buffer for 1 hour on rolling as previously published ^56^. After incubation, the suspension was collected and centrifugated at 4,000 g at 4°C for 4 minutes to pellet any large particles or debris. Total viruses in dust suspension were concentrated using polyethylene glycol (PEG) similar to virus concentration in sewage published previously ^57,58^. Briefly, PEG 6000 was added to each dust suspension to make up a final 10% PEG 6000 (Sigma-Aldrich, USA) concentration, followed by pH adjustment to pH 4 and overnight incubation at 4°C with shaking. After overnight incubation, sample was centrifuged at 13,500 g at 4°C for 90 minutes. Supernatant was removed and the remaining pellet was resuspended in 500 µL of pre-warmed glycine buffer, and then subjected to 5-minute centrifugation at 13,000 g at 4°C. Supernatant was collected, and supernatants from EDC samples that were collected in the same farm at the same time point (N=1-4) were pooled together for further processing. Viral-enriched dust suspension samples were then treated with TURBO DNase as previously described ^55^ to remove non-encapsulated DNA, followed by total nucleic acid extraction using MagMAX™ viral RNA isolation kit (Thermo Fisher, USA) according to manufacturer’ instructions but without the use of carrier RNA. Reverse transcription (using SuperScript III reverse transcriptase [Invitrogen, USA] and non-ribosomal hexamers ^59^) and second strand cDNA synthesis (using Klenow fragment 3’-5’ exo- [New England BioLabs, USA]) of chicken fecal samples and dust samples was performed as previously described ^55^ and following manufacturer’ instructions.

Standard Illumina libraries were prepared using Nextera XT DNA library kit following the manufacturers’ instructions. Final libraries were sequenced on the Illumina MiSeq platform (600 cycles; paired-end 2×300 bp). Chicken fecal samples were sequenced with a multiplex range of 16-19 samples per run. Dust samples and negative blank controls were sequenced with a multiplex of 12 samples per run.

### 4.3 Sequencing data analysis

Raw reads were subjected to adapter removal using Trim Galore/default Illumina software, followed by quality trimming using QUASR ^60^ with a threshold of minimum length of 125 nt and median Phred score ≤ 30. The resulting quality-controlled reads were *de novo* assembled using metaSPAdes v3.12.0 ^61^. *De novo* assembled contigs were classified using UBLAST v11.0667 ^62^ against eukaryotic virus family protein databases as previously described ^23,26^. We set a detection threshold of contig with minimum amino acid identity of 70%, minimum length of 300 nt and an e-value threshold of 1 × 10^−10^ when interpreting our contig classification results. Contig classification results were analyzed and visualized using R packages including dplyr, reshaped2 and ComplexHeatmap ^63–65^ and python package <u>squarify</u>.

### 4.4 Phylogenetic analysis

Sequences with ≥ 80% of genome coverage from the *Picornaviridae, Astroviridae, Caliciviridae* and *Parvoviridae* families were used for phylogenetic analyses, comparing them with global sequences retrieved from GenBank. Multiple sequence alignments were performed using MAFFT v7.427 ^66^ and manually checked in Geneious v2021.0.3 (Biomatters, New Zealand). Best-fit models of evolution for phylogenetic tree reconstruction were estimated using ModelFinder module in IQ-TREE v1.6.11 ^67^, determined by Akaike Information Criterion (AIC). Maximum-likelihood trees were constructed using RAxML-NG v1.0.1 ^68^ with 100 pseudo-replicates. Resulting trees were visualized and annotated using R package ggtree ^69^.

### 4.5 Kmer analysis with MASH

To compare Jaccard distances between dust samples, all quality controlled short reads, or resulting assembled viral contigs were analyzed using the Triangle function from Mash v2.3 ^43,44^ with a kmer size of 32 (-k 32) and a sketch size of 10,000 (-s 10000). The resulting distances were visualized in a heatmap using R package ggplot2 ^70^.

### 4.6 Data availability

The raw reads are available in the SRA under the BioProject accession number <u>PRJNA670873</u> (chicken feces) and <u>PRJNA701384</u> (farm dust samples) (**Table S1**). All sequences in phylogenetic analysis have been deposited in GenBank under the accession numbers MW684778 to MW684847 (**Table S2**).

## ACKNOWLEDGMENTS

We thank all the participating farms for their cooperation, Pella van der Wal and Gaby van Dijk from Erasmus MC for providing sequencing service. This virome study was funded by ZonMW TOP project 91217040 (K.T.T.K., M.M.T.dR., L.A.M.S., D.J.J.H. and M.P.G.K.) and a Marie Sklodowska-Curie Individual Fellowship, supported by Horizon 2020 (grant no. 799417; M.V.T.P.). M.V.T.P and M.C. was additionally supported by the Wellcome Trust and FCDO (grant agreement number 220977/Z/20/Z, awarded to M.C.). Sample collection was supported with internal funds from the Utrecht University (M.M.T.dR., A.B.M., and I.M.W.).

## AUTHOR CONTRIBUTION

K.T.T.K., M.P.G.K., M.C. and M.V.T.P. conceived the study. K.T.T.K. and M.V.T.P. performed experiments. M.M.T.dR and I.M.W. designed sampling strategy. M.M.T.dR. and A.B.M. conducted fieldwork, handled sample collection and collected associated metadata. K.T.T.K., M.C. and M.V.T.P. analyzed data. K.T.T.K. drafted the manuscript. K.T.T.K., M.V.T.P., M.C., M.P.G.K., L.A.M.S., I.M.W., M.M.T.dR and A.B.M. edited the manuscript. M.V.T.P., M.P.G.K. and I.M.W. supervised the study. M.P.G.K., M.C., D.J.J.H. and L.A.M.S. acquired the funding.

## Figures

**Figure S1.**
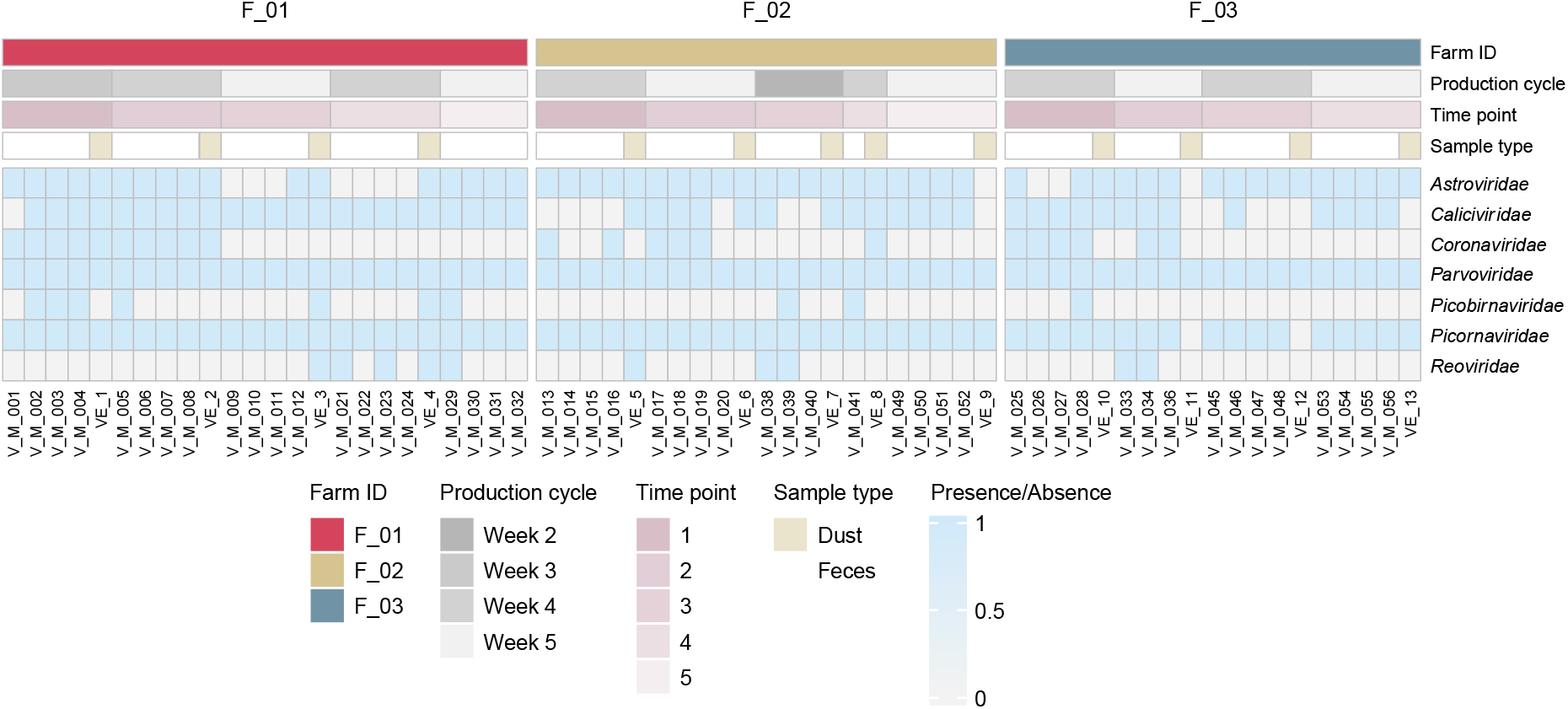
Detection of viral sequences from different virus families in chicken feces and farm dust samples. Associated metadata is shown at the top panel. The color intensity of the heatmap (bottom panel) is determined by number of contigs with minimum length of 300 nt, at least 70% identity and an e-value threshold of 1×10^−10^ when comparing with closest reference in our database. Sample order is assorted by farm ID, sampling time point. Time point is an arbitrary number. Samples were collected at the same time and place when both Farm ID and Time point match.

**Figure S2.**
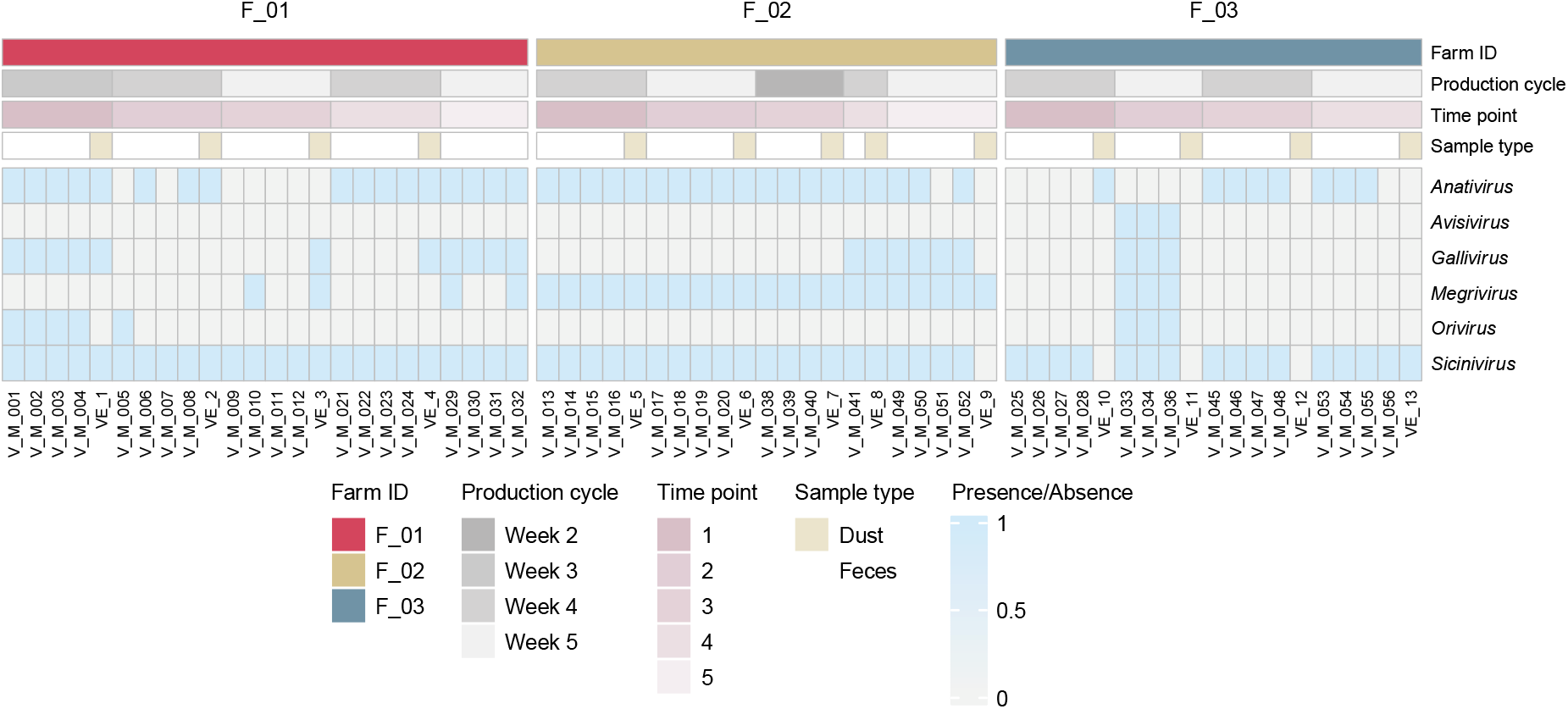
Detection of viral sequences from picornavirus genera in chicken feces and farm dust samples.Associated metadata is shown at the top panel. The color intensity of the heatmap (bottom panel) is determined by number of contigs with minimum length of 300 nt, at least 70% identity and an e-value threshold of 1×10^−10^ when comparing with closest reference in our database. Sample order is assorted by farm ID, sampling time point. Time point is an arbitrary number. Samples were collected at the same time and place when both Farm ID and Time point match.

**Table S1.** A list of corresponding BioProject, BioSample and SRA accession numbers for all samples.

**Table S2.** A list of corresponding GenBank accession numbers for sequences used in phylogenetic analyses.

